# Prioritization of Lung Cancer Candidate Genes using Moment of Inertia Tensor Analysis

**DOI:** 10.1101/2024.10.18.619023

**Authors:** Ayushi Dwivedi, Mallikarjuna Thippana, P. Manimaran, Vaibhav Vindal

**Affiliations:** Department of Biotechnology and Bioinformatics, School of Life Sciences, University of Hyderabad, Gachibowli, Hyderabad 500046, India; School of Physics, University of Hyderabad, Gachibowli, Hyderabad 500046, India

**Keywords:** Lung cancer, Candidate genes, Sequence similarity, Alignment-free method, Moment of Inertia, Survival analysis

## Abstract

A variety of factors contribute to the complexity of lung cancer progression. To comprehend the disease, candidate genes must be investigated. Present study aimed to employ an alignment-free method to prioritize candidate genes based on the physicochemical properties of the amino acids. It uses the moment of inertia tensor that measures the mass distribution around an axis of rotation, to compute the rotational energy and angular momentum of amino acids in protein sequences. The computed features were compared to those of established lung cancer genes, leading to the identification of 26 candidate genes with a high degree of similarity. These genes participate in critical biological processes that regulate the mitotic cell cycle and cell development. The prognostic significance of these genes was also assessed and four genes (IL1A, CDC25C, IL4R, and TGFBR1) were found to be associated with poor survival. Additionally, the role of prioritized genes and potential drugs that target these genes in other cancer types was also examined. Our method will help to discover new biomarkers and intervention strategies for lung cancer.

## 1. Introduction

Lung cancer is a prevalent and deadly type of cancer worldwide, characterized by uncontrolled cell growth in lung tissues. It affects millions of individuals annually and is responsible for a significant number of cancer-related mortalities due to its aggressive nature and the difficulty in early detection. Lung cancer is primarily classified into two main categories: small-cell lung cancer (SCLC) and non-small cell lung cancer (NSCLC). SCLC accounts for approximately 15-20% of all lung cancer cases, while NSCLC accounts for the remaining 80-85%. NSCLC has three subtypes: adenocarcinoma (LUAD, about 40%), squamous cell carcinoma (LUSC, around 25-30%), and large cell carcinoma (roughly 10-15%) [1–3]. The most significant risk factor for lung cancer is smoking; other risk factors include age, radon exposure, environmental pollution, occupational exposure, gender, race, and pre-existing lung diseases [4]. Despite these risk factors, recent advancements in tumor biology, prognostic genes, and treatment options have improved patient outcomes [5]. Public health interventions aimed at reducing smoking have also led to decreased lung cancer rates and increased survival rates in some countries. However, low-income countries still face a high burden of lung cancer due to limited access to healthcare and ineffective tobacco control policies [6–7].

Various computational methods have facilitated the prediction of candidate genes, categorized into exploring sequence similarities with known disease genes or assessing gene functional annotation [8–9]. Commonly used methods include ENDEAVOUR, G2D, SUSPECTS, GFSST, and POCUS [10–14]. However, experimental validation is resource-intensive and impractical for all predictions, necessitating the selection of the most promising candidates using reliable criteria [15]. Despite numerous studies identifying candidate genes for lung cancer, current bioinformatics tools inadequately prioritize them. Thus, developing a method to effectively prune and rank genes for further evaluation is essential. The most common methods for investigating sequence similarities among proteins are alignment-based, aiding in understanding functionally similar proteins [15–23]. However, alignment-free methods, such as the moment of inertia tensor originally developed for DNA sequences, offer advantages in comparison to alignment-based methods [24–25]. The moment of inertia tensor has proven efficient and reliable in studying the similarities among the protein sequences. Later, Hou et al., implemented the same method to analyze the sequence similarities between the proteins [26]. Recently, Thummadi et al. used the moment of inertia tensor to predict candidate genes associated with cervical cancer progression [27].

This study focuses on prioritizing candidate genes associated with lung cancer by employing the moment of inertia tensor concept. The moment of inertia tensor could serve as a component of a more complex method that analyzes protein structure and dynamics in addition to sequence. This approach quantifies the distribution of mass within a three-dimensional object, calculating the protein’s resistance to rotation around different axes. It enables the calculation of similarities between sequences of known lung cancer genes (KLCs) and candidate lung cancer genes (CLCs) in an alignment-free manner. This methodology not only optimizes resource utilization but also minimizes the effort required to prioritize candidate genes. Additionally, a comprehensive analysis was conducted to correlate the predicted candidate genes with relevant Gene Ontology (GO) terms and KEGG pathways, with the aim of elucidating their potential roles in diverse cancer types. Moreover, drug targets associated with these genes were systematically compiled. Through this comprehensive investigation, a prioritized list of genes and potential biomarkers pertinent to lung cancer was identified.

## 2. Materials and Methods

### 2.1 Data Curation

A total of 702 genes associated with lung cancer progression were systematically curated from the MalaCards database. Additionally, the Network of Cancer Genes (NCG7.0) database was utilized to identify 591 known and 2756 putative cancer-related genes [28]. Comparative analysis of these datasets yielded a shared set of 301 genes, categorized into 170 (KLC) genes and 131 (CLC) genes. To facilitate subsequent tensor analysis, protein sequences corresponding to the 301 lung cancer genes were acquired and processed using the R programming environment [29–31].

### 2.2 Three-dimensional model representation of protein sequence

In bioinformatics, amino acid sequence arrangement is crucial for determining protein three-dimensional structure, which in turn defines its functionality. Proteins are essentially composed of long chains of amino acid residues. Each protein is made up of twenty different amino acids, which determine its various functions. Recently, Hou et al. [26] developed a graphical representation method to visualize protein sequences as a three-dimensional model. The approach utilizes the physicochemical properties of each amino acid, such as hydrophobicity and molecular mass, as descriptors. The twenty amino acids are divided into two groups: hydrophilic (HPB) and hydrophobic (HFP) amino acids. HPB includes amino acids [E, G, D, N, K, H, Q, R, S, T] and HFP includes [C, A, V, P, F, L, M, Y, I, W]. Furthermore, the two groups of amino acids were sub-grouped based on their strong or weak strength. Hydrophilic amino acids that are weak denoted with WP = [H, Q, E, T, G]; and strong with SP = [K, D, S, R, N]; likewise, Hydrophobic amino acids that are weak denoted with WH = [V, P, M, C, A]; strong with SH = [W, Y, L, I, F]. To facilitate the calculation of the moment of inertia tensor, a simplified spatial representation of the protein sequence is employed. This representation assumes the amino acids are positioned sequentially along the circumference of a circle with a radius of one unit, with each hydrophobic amino acid in the first two quadrants and each hydrophilic amino acid in the third and fourth quadrants. Amino acid residues are arranged in ascending order within each quadrant, with abbreviated symbols representing each amino acid. Every amino acid is referred to with a coordinate (α_i_, β_i_), i.e., α_i_ = cos (2πi/20), and β_i_ = sin(2πi/20), where i = 1, 2 20. The γ axis coordinate values were obtained from the residual weight of each amino acids. The value +1 was assigned to heavier amino acids, and -1 was assigned to smaller amino acids for γ coordinates **(Table 1)**.

**Table 1.** Coordinates (α, β, γ) of amino acids and their residue weight.

| | Amino acid | Abbreviation | $\alpha$ | $\beta$ | $\gamma$ | Residue weight |
| --- | --- | --- | --- | --- | --- | --- |
| **Quadrant 1**<br>**Weak**<br>**Hydrophobic** | Alanine | Ala | 0.9511 | 0.3090 | -1 | 71.8 |
|  | Cysteine | Cys | 0.8090 | 0.5878 | -1 | 103.14 |
|  | Methionine | Met | 0.5878 | 0.8090 | 1 | 131.19 |
|  | Proline | Pro | 0.3090 | 0.9511 | -1 | 97.12 |
|  | Valine | Val | 0.0000 | 1.0000 | -1 | 99.13 |
| **Quadrant 2**<br>**Strong**<br>**Hydrophobic** | Phenylalanine | Phe | -0.3090 | 0.9511 | 1 | 147.17 |
|  | Isoleucine | Ile | -0.5878 | 0.8090 | -1 | 113.16 |
|  | Leucine | Leu | -0.8090 | 0.5878 | -1 | 113.16 |
|  | Tryptophan | Trp | -0.9511 | 0.3090 | 1 | 186.21 |
|  | Tyrosine | Tyr | -1.0000 | 0.0000 | 1 | 163.18 |
| **Quadrant 3**<br>**Strong**<br>**Hydrophilic** | Aspartic acid | Asp | -0.9511 | -0.3090 | 1 | 115.09 |
|  | Lysine | Lys | -0.8090 | -0.5878 | 1 | 128.17 |
|  | Asparagine | Asn | -0.5878 | -0.8090 | -1 | 114.1 |
|  | Arginine | Arg | -0.3090 | -0.9511 | 1 | 156.19 |
|  | Serine | Ser | 0.0000 | -1.0000 | -1 | 87.08 |
| **Quadrant 4**<br>**Weak**<br>**Hydrophilic** | Glutamic acid | Glu | 0.3090 | -0.9511 | 1 | 129.12 |
|  | Glycine | Gly | 0.5878 | -0.8090 | -1 | 57.05 |
|  | Histidine | His | 0.8090 | -0.5878 | 1 | 137.14 |
|  | Glutamine | Gln | 0.9511 | -0.3090 | 1 | 128.13 |
|  | Threonine | Thr | 1.0000 | 0.0000 | -1 | 101.11 |

The 3D protein sequence models were constructed using 3D Cartesian coordinates assigned to the amino acids. The centre of mass (m) of the 3D Cartesian coordinate system is given as,

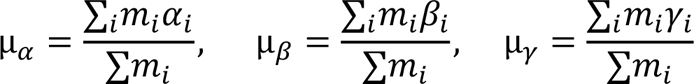

where α_i_, β_i_, γ_i_ denotes the *m_i_* coordinates. The moment of inertia tensor is denoted as a matrix,

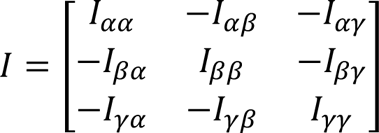

The components of the inertia matrix abbreviated as,

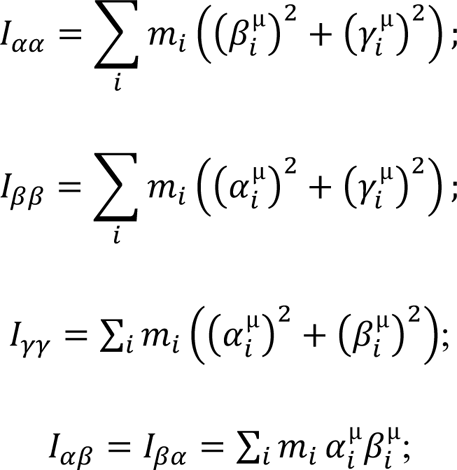

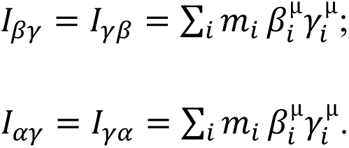

where 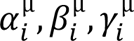 represents the Cartesian coordinates of the centre of mass *m_i_* which is the origin of the system. *λ*_1_, *λ*_2_, *λ*_3_ are the eigenvalues of the inertia matrix ‘I’ and protein sequence can be defined by a vector form represented as 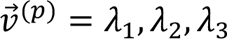 and Euclidean distance D is used to calculate the similarity score among the two protein sequences (P^1^, P^2^). Hence, distance 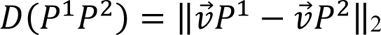. This indicates that the similarity score of proteins is inversely proportional to the distance between the two proteins. Therefore, this method has the potential to observe similarities among sequences of having different lengths [26].

### 2.3 Functional enrichment analysis

The investigation of a prioritized list of candidate genes was carried out to determine their association with significant GO terms and KEGG pathways using functional enrichment analysis. The biological processes (BP), molecular functions (MF), and cellular components (CC) of each gene was examined in relation to GO terms, while pathways were analyzed using the KEGG pathway database. The clusterProfiler R package [32] was employed to interpret the functional roles of the candidate genes.

### 2.4 Survival analysis

For the prioritized candidate genes, survival analysis was performed to validate the effect of the gene expression pattern on patient survival using the Kaplan-Meier (KM) Plotter [33]. Survival analysis provides valuable insights into time-to-death events, and our results indicate that a higher or lower level of gene expression pattern is associated with poor survival outcomes in lung cancer patients.

## 3. Results

### 3.1 Lung cancer gene categorization

The gene expression profiles of 170 (KLC) and 131 (CLC) lung cancer-associated genes were analyzed using the GEPIA2 web server. [34]. This tool analyzes gene expression profiling based on data from the TCGA (The Cancer Genome Atlas) [35]. The TCGA provides lung cancer gene expression data for two projects: LUAD and LUSC, which are differentiated based on the location of the tumor’s origin.

Among the 170 KLC genes, 62 genes were found to be overexpressed or upregulated, while 108 genes were downregulated in patients with LUAD. In the case of LUSC patients, 79 genes were upregulated, and 91 genes were downregulated. Among the 131 CLC genes, 60 were upregulated and 71 were downregulated in LUAD patients. For LUSC patients, 57 genes were upregulated and 74 were downregulated. The bar plot depicts the distribution of the genes [**Fig. 1**].

**Fig. 1:**
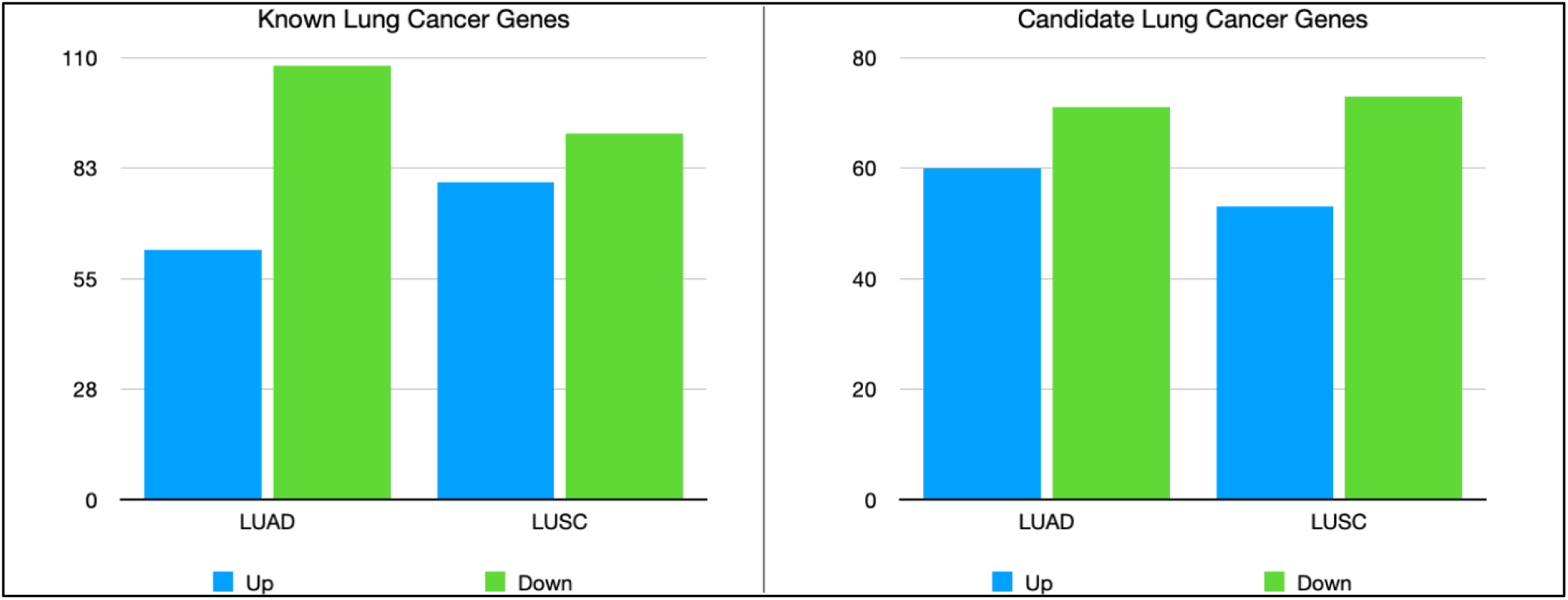
Number of upregulated and downregulated genes from known and candidate gene list

### 3.2 Tensor analysis on the known disease genes and candidate genes

The moment of inertia tensor analysis on KLC and CLC proteins identifies the closeness of the respective proteins. The Euclidean distance between the KLC and CLC genes was calculated from the moment of inertia matrix. The distance is calculated based on the largest Eigenvalue of each protein sequence. The resulting matrix ranges from 2.213349 to 28892.520172. A dendrogram was constructed to visualize protein similarity, clustering proteins based on shared characteristics. (**Fig. 2**).

**Fig. 2:**
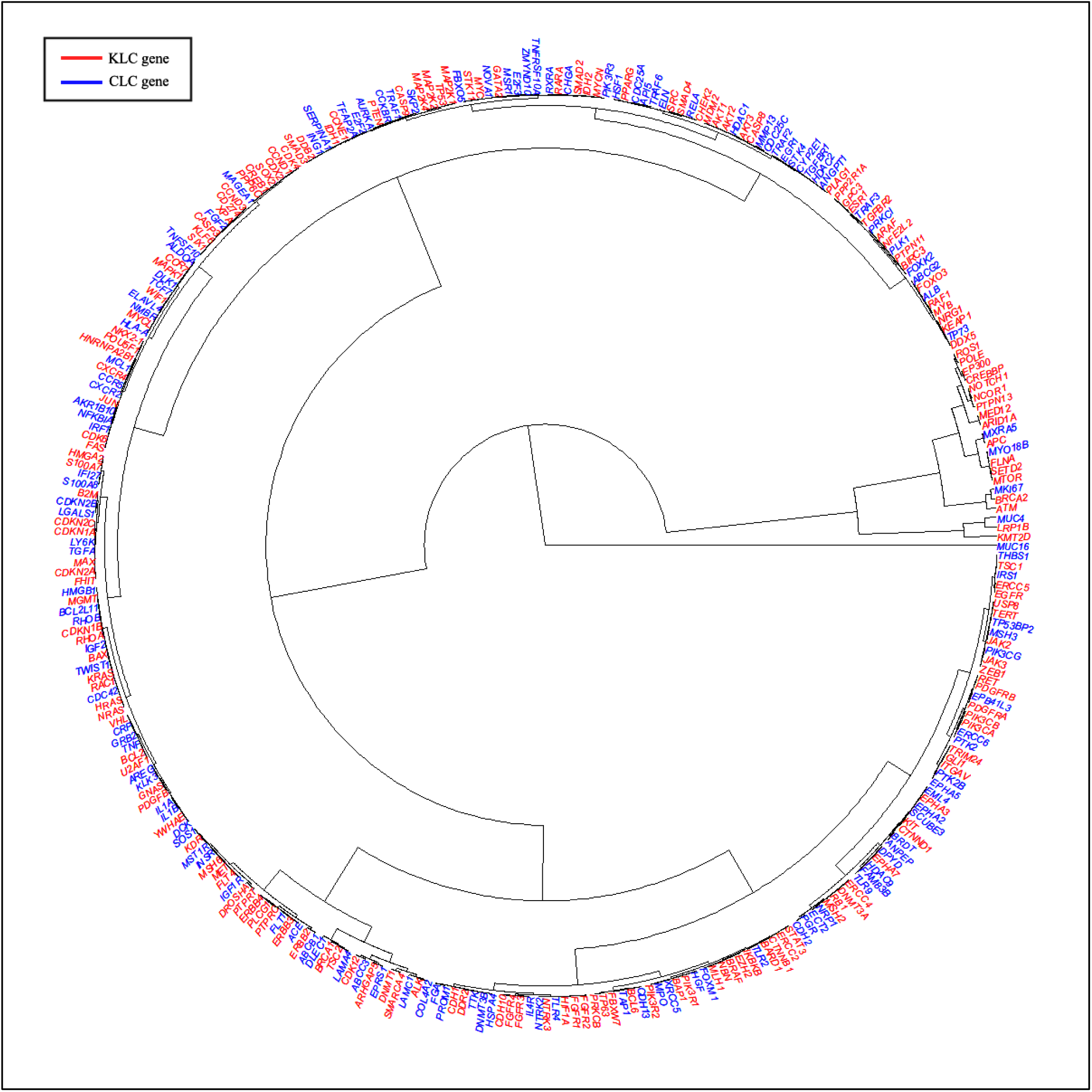
Circular dendrogram of known and candidate genes belong to lung cancer. Known lung cancer genes represented in red color and candidate lung cancer genes represented in blue color

Less similar proteins were MUC16 and CDKN2B based on the maximum distance. Employing the Euclidian distance matrix, CLC proteins were prioritized that have 5% or lower distance (>=95% similarity) to KLC proteins were compared to the larger distance (least similar) proteins. Among the CLC proteins, namely KMT2D, PLCG1, PLAG1, MKI67, TWIST1, ZMYND10 and AKR1B10 were found to have 95% similarity score with at least one or two maximum distance proteins from the KLC list. Subsequently, proteins exhibiting substantial similarity to multiple KLC proteins were prioritized, resulting in a refined set of 68 candidate proteins based on a number of associations with KLC. A stringent inclusion criterion of at least nine associations with KLC proteins was applied, yielding 26 candidate proteins previously unreported in lung cancer. The Therapeutic Target Database was referred to acquire comprehensive disease and drug-related information for these 26 proteins. **(Table 2)**.

**Table 2:**
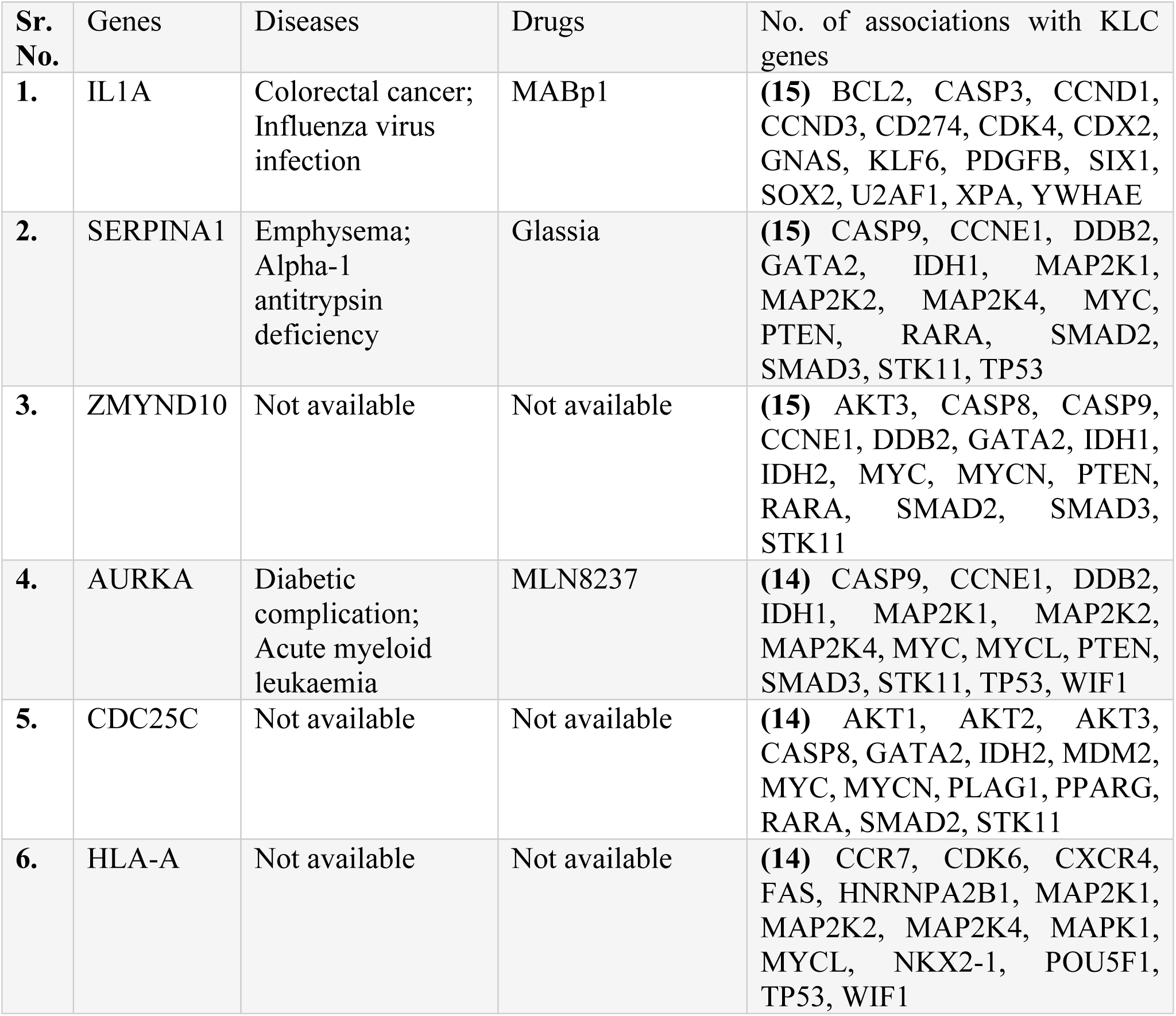
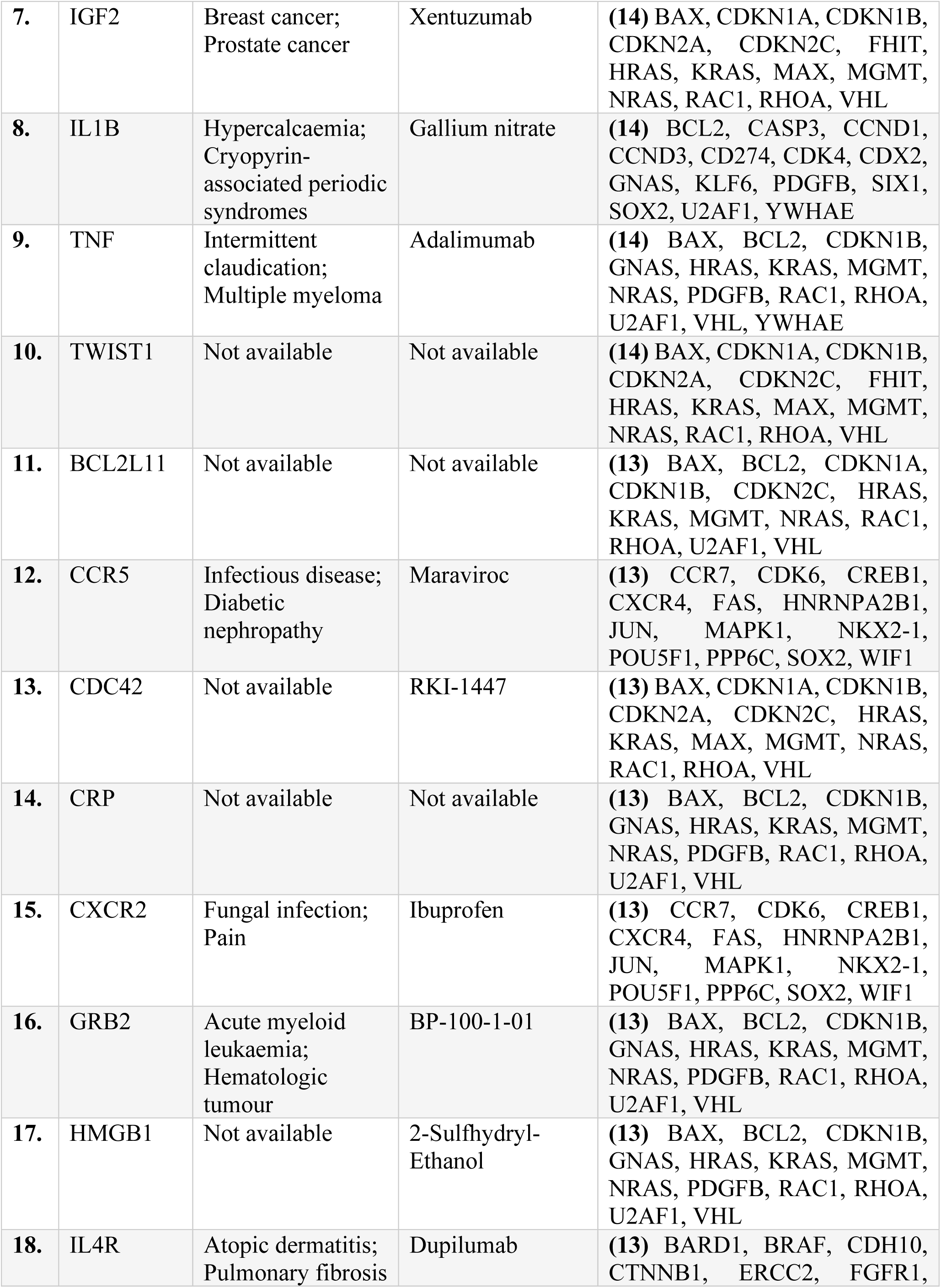
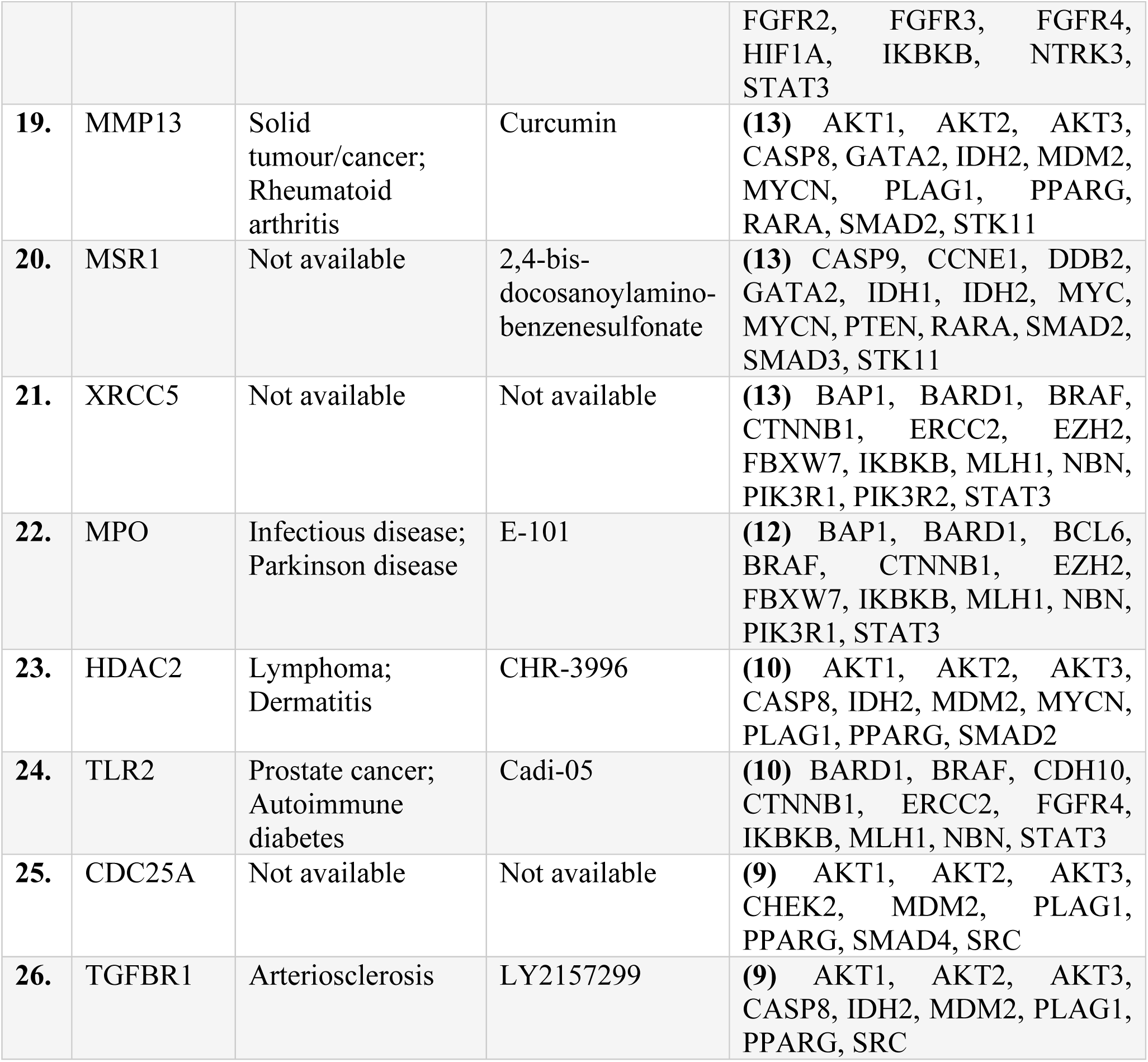
Prioritized biomarkers for the lung cancer.

### 3.3 Functional interpretation of potential candidate proteins

Functional interpretation of genes is imperative for understanding their molecular functions, biological processes, cellular components, and pathways. Functional enrichment analysis was performed on known lung cancer proteins and prioritized candidates. Terms exhibiting p-values less than 0.05 were considered.

#### Biological process

Known lung cancer genes were associated with significant GO terms of biological processes related to the cell cycle regulation. **(Table 3)**.

**Table 3:** Top 10 GO terms from biological processes.

| Sr. No. | GO: ID | GO: TERM | P-value | No. of genes |
| --- | --- | --- | --- | --- |
| 1. | GO:0042110 | T cell activation | 1.16E-08 | 9 |
| 2. | GO:0050863 | regulation of T cell activation | 2.91E-07 | 7 |
| 3. | GO:0050870 | positive regulation of T cell activation | 4.57E-07 | 6 |
| 4. | GO:1903039 | positive regulation of leukocyte cell-cell adhesion | 8.14E-07 | 6 |
| 5. | GO:0030099 | myeloid cell differentiation | 1.54E-06 | 7 |
| 6. | GO:0045785 | positive regulation of cell adhesion | 1.69E-06 | 7 |
| 7. | GO:0001819 | positive regulation of cytokine production | 2.03E-06 | 7 |
| 8. | GO:0022409 | positive regulation of cell-cell adhesion | 2.13E-06 | 6 |
| 9. | GO:0048145 | regulation of fibroblast proliferation | 4.65E-06 | 4 |
| 10. | GO:0048144 | fibroblast proliferation | 4.89E-06 | 4 |

For the prioritized genes, various significant biological processes were associated with, including the positive regulation of chemokine production, positive regulation of interleukin-6 production, regulation of chemokine production, chemokine production, acute-phase response, positive regulation of cell cycle, positive regulation of cytokine production, regulation of mitotic nuclear division, acute inflammatory response, positive regulation of cell development, regulation of mitotic cell cycle, and so on. **[Fig. 3]**.

**Fig. 3:**
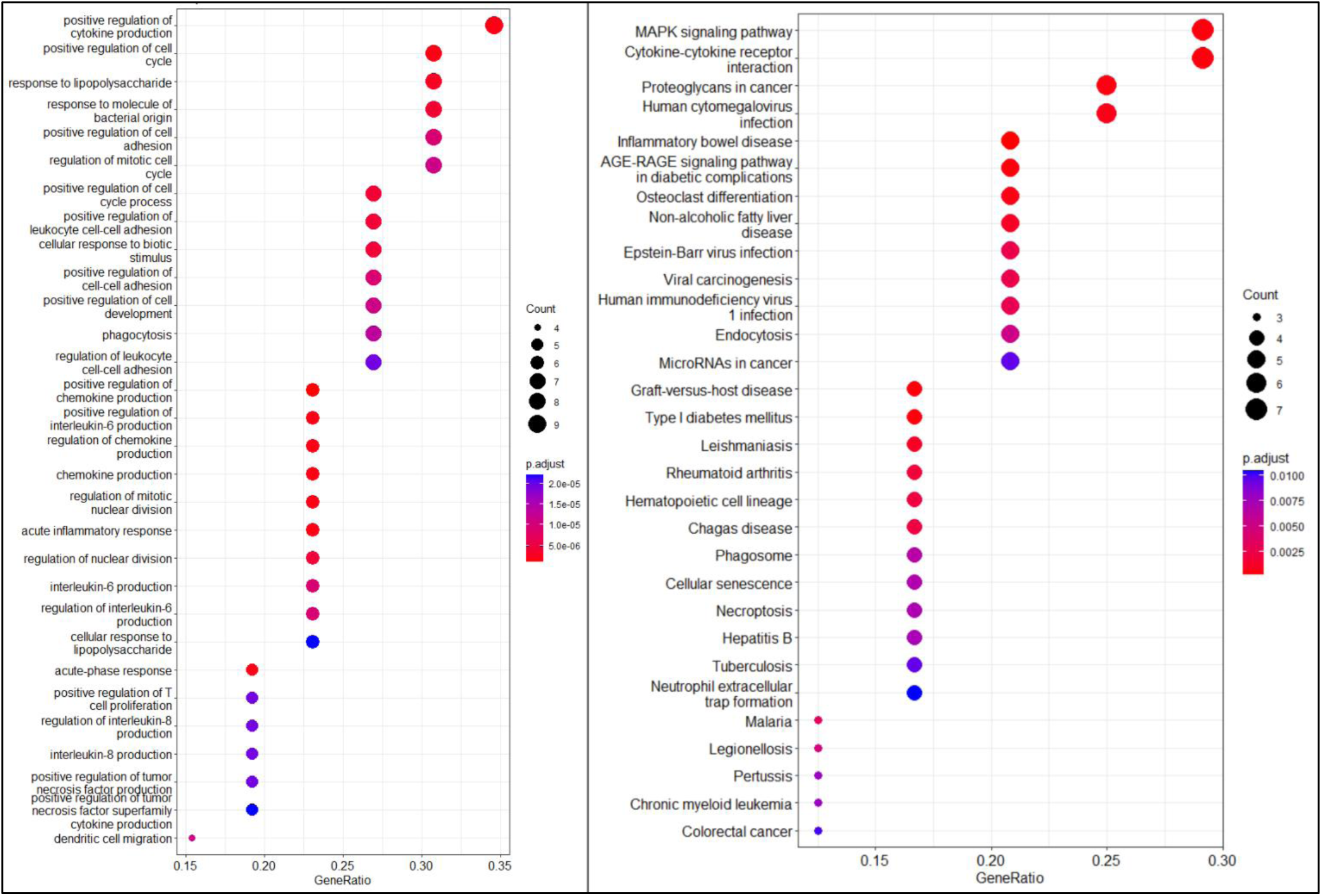
Functional interpretation of candidate genes for their biological relevance: Left panel shows GO terms: Biological processes and KEGG pathways in right panel.

#### KEGG Pathway analysis

These results indicate that the known lung cancer genes are associated with various cancer-related pathways. These enriched pathways are represented in tabular form **(Table 4).**

**Table 4:** Top 10 enriched KEGG pathways.

| Sr. No. | ID | KEGG Pathway | P-value | No. of genes |
| --- | --- | --- | --- | --- |
| 1. | hsa05167 | Kaposi sarcoma-associated herpesvirus infection | 1.16E-12 | 11 |
| 2. | hsa05163 | Human cytomegalovirus infection | 1.84E-10 | 10 |
| 3. | hsa05212 | Pancreatic cancer | 1.04E-09 | 7 |
| 4. | hsa05169 | Epstein-Barr virus infection | 1.86E-09 | 9 |
| 5. | hsa05203 | Viral carcinogenesis | 2.03E-09 | 9 |
| 6. | hsa05220 | Chronic myeloid leukemia | 4.92E-08 | 6 |
| 7. | hsa05162 | Measles | 1.65E-06 | 7 |
| 8. | hsa04151 | PI3K-Akt signaling pathway | 2.46E-07 | 8 |
| 9. | hsa05165 | Human papillomavirus infection | 1.38E-07 | 9 |
| 10. | hsa05160 | Hepatitis C | 1.68E-07 | 7 |

Inflammatory bowel disease, Graft-versus-host disease, Type I diabetes mellitus, AGE-RAGE signaling pathway in diabetic complications, the MAPK signaling pathway, Cytokine-cytokine receptor interaction, Proteoglycans in cancer, Osteoclast differentiation, Rheumatoid arthritis, Epstein-Barr virus infection, Viral carcinogenesis, Human immunodeficiency virus 1 infection, Pertussis, among other conditions, are influenced significantly by genes that are prioritized [**Fig. 3]**.

### 3.4 Survival analysis

The prognostic performance of candidate genes in patients with lung cancer was assessed using the Kaplan–Meier survival curve and log-rank test. Genes with poor prognosis were filtered based on a hazard ratio >1 and log-rank p-value <0.05. A total of 26 genes were prioritized for their poor prognosis in lung cancer patients, and this information is provided in a tabular format **(Table 5)**, i.e., 11 genes in LUAD and 9 genes in LUSC **(Fig. 4a and 4b)**

**Table 5:** List of prioritized genes in lung cancer with information related to expression and survival analysis.

| Sr. No. | Genes | Description | LUAD<br>(Expression pattern)<br>(Upregulated/<br>Downregulated) | LUSC<br>(Expression pattern)<br>(Upregulated/<br>Downregulated) | Survival<br>Analysis<br>Result<br>(LUAD) | Survival<br>Analysis<br>Result<br>(LUSC) |
| --- | --- | --- | --- | --- | --- | --- |
| 1. | IL1A | Interleukin-1 alpha | Downregulated | Upregulated | Yes | Yes |
| 2 | SERPINA1 | Alpha-1-antitrypsin | Downregulated | Downregulated | No | Yes |
| 3 | ZMYND10 | Zinc Finger MYND-Type Containing 10 | Downregulated | Downregulated | No | No |
| 4 | AURKA | Aurora kinase A | Upregulated | Upregulated | Yes | No |
| 5 | CDC25C | Cell Division Cycle 25C | Upregulated | Upregulated | Yes | Yes |
| 6 | HLA-A | MHC class I antigen A*33 (HLA-A) | Upregulated | Downregulated | No | No |
| 7 | IGF2 | Insulin-like growth factor-II | Downregulated | Downregulated | No | No |
| 8 | IL1B | Interleukin-1 beta | Downregulated | Downregulated | No | Yes |
| 9 | TNF | Tumor necrosis factor | Downregulated | Downregulated | No | Yes |
| 10 | TWIST1 | Twist-related protein 1 | Upregulated | Upregulated | Yes | No |
| 11 | BCL2L11 | BCL2 Like 11 | Upregulated | Upregulated | No | No |
| 12 | CCR5 | C-C chemokine receptor type 5 | Downregulated | Downregulated | No | No |
| 13 | CDC42 | Cell Division Cycle 42 | Downregulated | Downregulated | No | No |
| 14 | CRP | HUMAN c-reactive protein | Downregulated | Downregulated | Yes | No |
| 15 | CXCR2 | C-X-C chemokine receptor type 2 | Downregulated | Downregulated | No | No |
| 16 | GRB2 | Growth Factor Receptor Bound Protein 2 | Downregulated | Downregulated | No | Yes |
| 17 | HMGB1 | High Mobility Group Box 1 | Downregulated | Upregulated | Yes | No |
| 18 | IL4R | Interleukin 4 receptor alpha | Downregulated | Downregulated | Yes | Yes |
| 19 | MMP13 | Matrix metalloproteinase-13 | Upregulated | Upregulated | No | No |
| 20 | MSR1 | Macrophage Scavenger Receptor 1 | Downregulated | Downregulated | No | Yes |
| 21 | XRCC5 | X-ray repair cross-complementing 5 | Upregulated | Upregulated | Yes | No |
| 22 | MPO | Myeloperoxidase | Downregulated | Downregulated | No | No |
| 23 | HDAC2 | Histone deacetylase 2 | Upregulated | Upregulated | Yes | No |
| 24 | TLR2 | Toll-like receptor 2 | Downregulated | Downregulated | No | No |
| 25 | CDC25A | Cell Division<br>Cycle 25A | Upregulated | Upregulated | Yes | No |
| 26 | TGFBR1 | Transforming<br>Growth Factor<br>Beta Receptor 1 | Downregulated | Downregulated | Yes | Yes |

**Fig. 4a:**
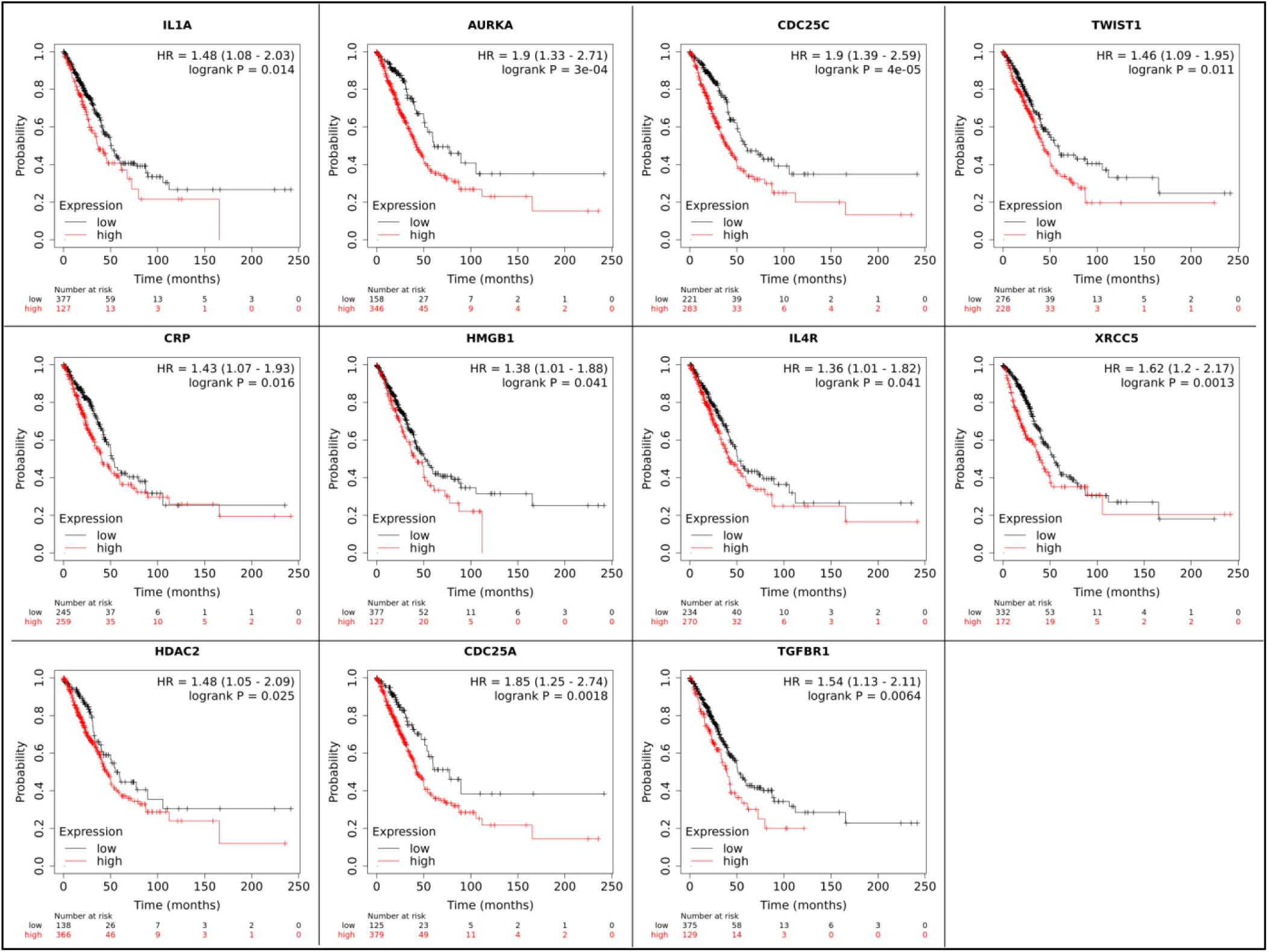
Prioritized genes that showed poor prognosis in LUAD patients

**Fig. 4b:**
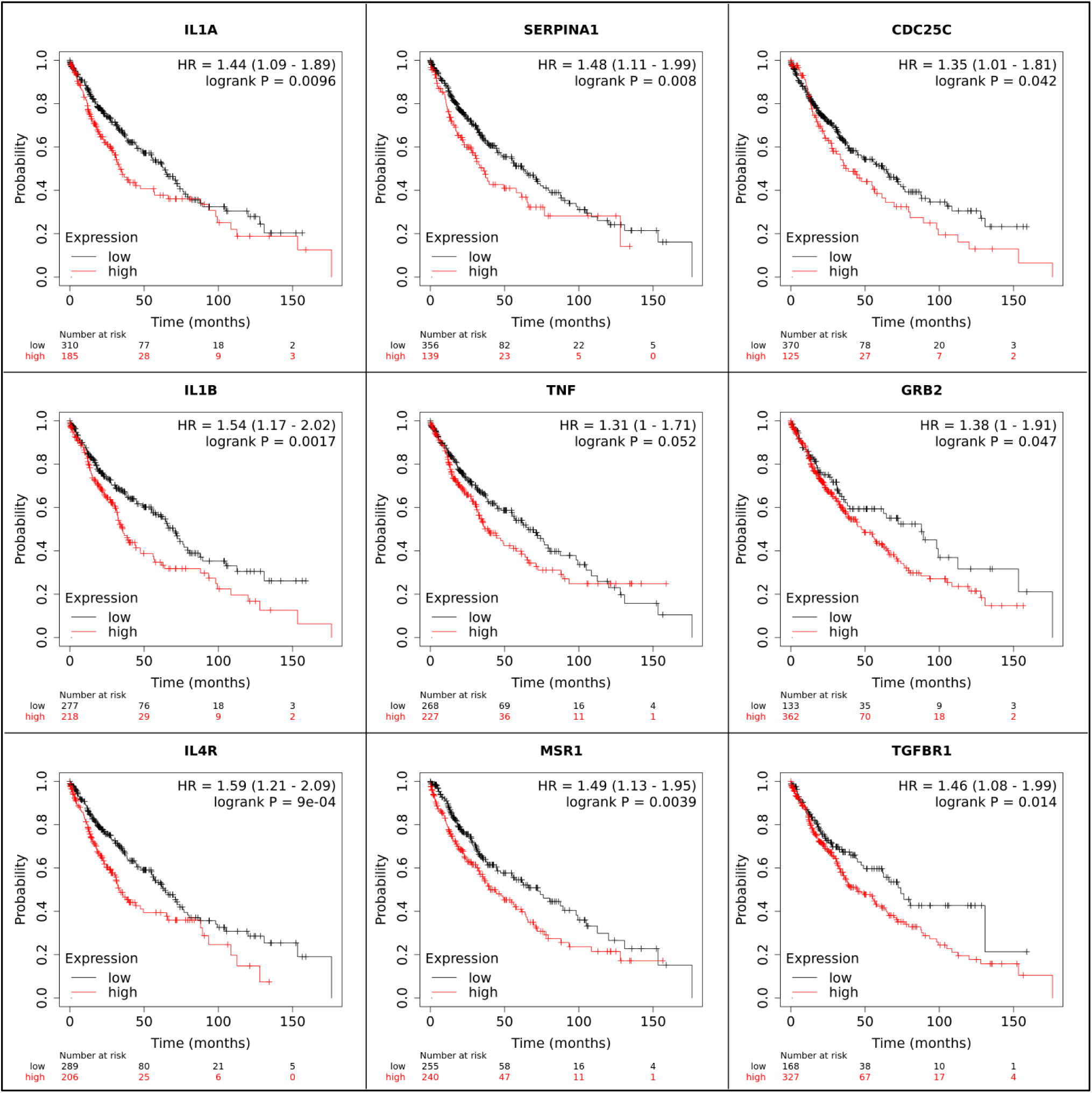
Prioritized genes that showed poor prognosis in LUSC patients

This study investigates the prognostic significance of four key genes in patients with both LUAD and LUSC. The expression of these genes, when downregulated or upregulated, can have detrimental effects on the prognosis of lung cancer patients.

## 4. Discussion

Lung cancer is known to have a higher mortality rate than all other cancers except for breast cancer. This study implemented an alignment-independent approach to examine protein sequences derived from established disease genes and those implicated in lung cancer tumorigenesis. The findings show the usefulness of tensors for the moment of inertia in prioritizing potential candidate proteins for lung cancer. Previous studies have reported prognostic/diagnostic biomarkers in lung carcinoma progression through various computational methods [36–37].

The aim of this study was to investigate sequence similarities among proteins using an alignment-free method. Specifically, the moment of inertia tensor was employed to compute distance metrics between proteins, encompassing established and putative genes, based on their physicochemical properties. In previous studies, the alignment-free method was effectively used to compare 12 baculovirus sequences with traditional clustalX-based alignment [26]. Our analysis revealed that the protein sequences with the highest similarity belonged to the same family, as determined through tensor analysis. Notably, pairs of highly similar proteins among KLC proteins belonging to the same family include TGFBR2 – ATM (serine/threonine protein kinase family), ALK – IGF1R (receptor tyrosine kinase family), ARAF – BRAF (RAF family), and HRAS – KRAS (RAS oncogenic family). Similarly, protein pairs with the highest similarity among CLC and KLC proteins were found to be associated with the same family, such as BCL2 – TNF and TGFA – CDKN2A. Tensor-based approaches have proven highly effective in discerning similarities among protein sequences. However, a limitation emerges in differentiating between sequences of identical length and amino acid composition but varying arrangement. Although such instances may be infrequent, they remain a potential challenge for proteins characterized by comparable amino acid content. This analysis revealed 26 genes not previously documented in lung cancer within the GeneCards database. Notably, among these novel candidates, IL1A, CDC25C, IL4R, and TGFBR1 demonstrated unfavorable prognostic indicators as assessed by Kaplan-Meier survival analysis.

Interleukin-1 (IL-1) is a cytokine signalling molecule that serves as a potent regulator of immune and inflammatory responses. It has a crucial role in carcinogenesis and tumor progression, particularly in neuroinflammation. The presence of elevated levels of TNF and IL-1 in the brain can lead to the breakdown of the blood-brain barrier, a critical component of the central nervous system [38–40]. Furthermore, mutations in IL-1 have been linked to an increased risk of developing gastric cancer, Graves’ disease, and ankylosing spondylitis [41–43]. This cytokine was found to be associated with 20 significant KEGG pathway enrichment terms, including inflammatory bowel disease, pertussis, leishmaniasis, and MAPK signaling pathways. In the GO enrichment analysis of biological processes, 113 significant terms were identified, highlighting associations such as fever generation, regulation of immature T cell proliferation, programmed cell death involved in cell development, inflammatory response to wounding, and positive regulation of tumor necrosis factor production.

Cell Division Cycle 25C (CDC25C) is a critical cell cycle regulatory protein that plays a vital role in regulating G2/M cell cycle progression and checkpoint proteins in DNA damage repair. Recently, research has reported that alterations in CDC25C expression levels are associated with tumorigenesis and tumor development, and it may be a potential therapeutic target for cancer treatment. Various studies have demonstrated that increased expression is associated with poor prognosis and low survival rates in lung, liver, gastric, bladder, colorectal, esophageal, and prostate cancers and acute myeloid leukemia [44]. Functional enrichment analysis found associations with five significant KEGG pathway terms, including progesterone-mediated oocyte maturation, cell cycle, oocyte meiosis, microRNAs in cancer, and human immunodeficiency virus 1 infection. Furthermore, GO enrichment of biological processes revealed 66 significantly enriched terms, such as positive regulation of G2/M transition of the mitotic cell cycle, meiotic cell cycle phase transition, and signal transduction involved in the mitotic G1 DNA damage checkpoint.

Interleukin-4 receptor (IL4R) is highly expressed in various types of malignancies, such as epithelial cancer metastasis [45]. It also contributes to the progression of malignant tumors, including breast, prostate, and epithelial cancers [46]. IL4R, in conjunction with other cytokines and macrophages, plays a crucial role in allergies by reducing the levels of pro-inflammatory cytokines during IgE synthesis [45]. IgE antibodies are vital for tumor control during natural tumor surveillance and in altered immunotherapy settings. It was associated with seven enriched terms in KEGG pathways, including inflammatory bowel disease, Th1 and Th2 cell differentiation, hematopoietic cell lineage, Th17 cell differentiation, JAK-STAT signaling pathway, cytokine-cytokine receptor interaction, and PI3K-Akt signaling pathways. Additionally, it was associated with enrichment in 141 GO terms of biological processes, including regulation of T-helper cell differentiation, positive regulation of myoblast fusion, and positive regulation of mast cell activation, which are involved in the immune response.

Transforming growth factor beta receptor type I (TGFBR1) belongs to the serine-threonine protein kinase family and has been associated with 22 significantly enriched KEGG pathway terms, such as adherens junctions, pancreatic cancer, chronic myeloid leukemia, colorectal cancer, TGF-beta signaling pathway, relaxin signaling pathway, FoxO signaling pathway, gastric cancer, Hippo signaling, and MAPK signaling pathways. Additionally, TGFBR1 is associated with 184 significant GO terms for biological processes, including endothelial cell activation, regulation of SMAD protein signal transduction, thymus development, and the activin receptor signaling pathway [47–48]. The vast majority of prioritized proteins have been extensively examined in a wide range of cancer types. Consequently, they are appealing candidates for exploration in the context of lung cancer and its development. Moreover, numerous drugs that target these proteins are currently available, which may prove to be effective treatments for lung cancer.

## 5. Conclusion

In conducted study, an alignment-free method based on the moment of inertia tensor was employed to investigate the sequence similarity between the KLC and CLC proteins. Potential biomarkers playing a crucial role in lung cancer progression were prioritized by exploring the sequence similarity between these proteins. This approach resulted in the identification of 26 genes that exhibited the highest degree of similarity to known lung cancer genes. Four genes, namely IL1A, CDC25C, IL4R, and TGFBR1, were proposed, which show poor prognosis in lung cancer patients as determined by survival analysis. These genes may be further investigated to develop novel prognostic biomarkers for the disease. Most of the prioritized genes have known associated drugs, which further enhances the significance of these identified proteins. A fast and efficient method was utilized to measure the sequence similarity between protein sequences, employing fewer parameters to describe the similarities between protein sequences, in contrast to other methods. The study highlights the utility of the moment of inertia tensor in ranking genes based on sequence similarity, and this method’s applicability to sequence similarity in other complex systems.

## Acknowledgment

The authors would like to thank CMSD, University of Hyderabad, for providing computational facilities. Vindal V would like to acknowledge the Institution of Eminence (IoE), University of Hyderabad (No. UoH/IoE/RC3-21-052), Indian Council of Medical Research (ICMR), GoI (ISRM/12(72)/2020, ID: 2020-2951), and Department of Biotechnology, GoI (No. BUILDER-DBT-BT/INF/22/SP41176/2020) for their financial support. Mallikarjuna T would like to thank ICMR, GoI, for the financial support as SRF (Ref. No: ISRM/11(47)/2019). Manimaran P would like to thank the Department of Science and Technology, Government of India, for their financial support (DST-SERB GoI Project No: YSS/2015/000949).

